# Parity and APOEε4 genotype contribute distinct changes to functional connectivity across the middle-aged brain

**DOI:** 10.1101/2025.03.01.640999

**Authors:** Bonnie H. Lee, Andrew J. McGovern, Stephanie E. Lieblich, Selene V. Mitchell, Jie Yin, Jonathan R. Epp, Liisa A.M. Galea

## Abstract

Cognition and its underlying neurobiology change throughout the trajectory of aging, with prominent sex differences and influences of sex-specific factors. Research has shown that parity (pregnancy and parenthood) uniquely altered various biomarkers of brain health in middle age depending on presence of Alzheimer’s disease (AD) risk. The present study builds on prior work by providing a comprehensive view of functional connectivity changes and elucidating how network-level dynamics contribute to cognitive outcomes depending on primiparity and APOEε4 genotype, the top genetic risk factor for late-onset sporadic AD risk.

We assessed neural activation in middle-aged wildtype and hAPOEε4 rats that were either nulliparous (0 litters) or primiparous (1 litter). Activation of the immediate early gene zif268 was quantified across 19 brain regions implicated in memory and AD. Primiparous hAPOEε4 rats exhibited widespread reductions in neural activation, particularly in the dorsal striatum, nucleus accumbens, frontal cortex, and retrosplenial cortex. Network analyses further revealed that primiparous wildtype rats had the most cohesive and efficient functional connectivity networks. Notably, the hierarchy of influence of brain regions within the neural network shifted based on parity and hAPOEε4 genotype. Activation of hippocampal new-born neurons in conjunction with subregions of the dorsal striatum, frontal cortex, and retrosplenial cortex dynamically predicted cognitive performance in a parity- and genotype-dependent manner. These findings underscore the lasting impact of reproductive history on brain health and cognitive aging, highlighting the need to consider sex-specific experiences in aging and AD research.

---

The trajectory of brain aging can be shaped by various factors, including genetic influences and lifestyle choices, and influences of sex are prominent across these relationships (Arenaza-Urquijo et al., 2024; Mattson & Arumugam, 2018; Qi et al., 2024; Roe et al., 2024; Subramaniapillai et al., 2021). Sex differences are seen in cognitive aging as well as in the neurobiology underlying cognitive aging (Bettio et al., 2017; Lee et al., 2023; Scahill et al., 2003). Significant hippocampal volume loss is seen throughout the process of aging, but how this varies by sex is equivocal. Although one study found faster decline in hippocampal volume in human females than males (Nobis et al., 2019), another study reported a later but faster decline in human males, and showed that age-related hippocampal volume decline was linear in females and quadratic in males in humans (Wang et al., 2019). It is important to underscore that sex differences in hippocampal volume can vary depending on the correction factor used to normalize for brain size differences (DeCasien et al., 2022; Duarte-Guterman et al., 2021; Sanchis-Segura et al., 2019) as well as other factors, including sex-specific experiences.

Beyond understanding sex differences in brain health, it is equally important to investigate how sex-specific experiences and factors may play a role. Indeed, previous parity (pregnancy and motherhood) is a critical modifier of brain and cognitive health, with long-lasting changes seen across aging (de Lange et al., 2020; Lindseth et al., 2022; Zhou et al., 2022). Research in humans has demonstrated a link between previous parity and reduced brain aging and slower cognitive decline in later life (de Lange et al., 2019; Zhou et al., 2022). Similarly, previous parity enhances hippocampus-dependent cognition and neuroplasticity at middle age in rodents (Barha & Galea, 2011; Duarte-Guterman et al., 2023; Eid et al., 2019; Galea et al., 2018; Kinsley et al., 1999). However, there is also evidence to suggest previous parity might contribute to increased risk of Alzheimer’s disease (AD) and AD-related biomarkers in some populations (Beeri et al., 2006; Colucci et al., 2006; Jang et al., 2018). These seemingly contrasting influences of parity on brain health have been referred to as the “parity paradox”, which postulates that parity might differentially influence biomarkers of brain health depending on presence of AD pathology or risk (Lee et al., 2024).

Late-onset sporadic AD makes up of over 95% of AD cases, and although the underlying causes remain unclear, there are known risk factors – the strongest of which include presence of Apolipoprotein E (APOE) ε4 alleles, advancing age, and female sex (‘2024 Alzheimer’s Disease Facts and Figures’, 2024; Riedel et al., 2016). Indeed, 60% of individuals with Alzheimer disease possess an APOE ε4 allele, and there is an interaction between female sex and APOEε4 genotype such that female carriers present with greater risk and burden of AD compared to male carriers (Duarte-Guterman et al., 2021; Farrer et al., 1997; Hohman et al., 2018; Koran et al., 2017). Previous research has examined the “parity paradox” in relation to cognition and various biomarkers of brain aging in rodent models of AD risk, including in humanized (h)APOEe4 rats (Cui et al., 2014; Lee et al., 2024). Previous parity in wildtype rodents is associated with improved cognitive performance and increased neuroplasticity (neurogenesis, synaptic proteins) at middle age (Barha et al., 2015; Cui et al., 2014; Galea et al., 2018; Lee et al., 2024; Zimberknopf et al., 2011). In contrast, in rodent models of AD risk, previous parity is associated with poorer cognitive performance, reduced hippocampal neurogenesis, and increased neuropathology in the hippocampus and cortex (Cui et al., 2014; Lee et al., 2024). Together, these findings suggest that previous parity alters the trajectory of brain aging and is an important factor to consider in understanding alterations in brain function with age and AD risk.

There has been considerable research to further our knowledge about functional connectivity changes related to AD pathogenesis, particularly in brain regions implicated in memory and reward processing, such as the hippocampus, frontal cortex, dorsal striatum, nucleus accumbens, and retrosplenial cortex (Grossman et al., 2003; Jobson et al., 2021; Perry & Kramer, 2015). These regions are among the earliest to exhibit connectivity disruptions and atrophy in AD (Cordella et al., 2018; Grossman et al., 2003; Nie et al., 2017; Terstege, Ren, et al., 2024). In individuals with mild cognitive impairment, a prodromal state to AD, whole brain metabolism was decreased in those who later developed AD compared to those who did not, and this effect was more prominent in human females compared to males (Terstege, Galea, et al., 2024). In addition, hypometabolism in the retrosplenial cortex was predictive of conversion to AD independent of amyloid or tau pathology, highlighting the retrosplenial cortex as an early site of metabolic dysfunction in subjects at risk for developing AD (Terstege, Galea, et al., 2024). In AD, impaired dynamics of excitatory and inhibitory synaptic function are evident across diverse neuronal subpopulations and are associated with deposition of amyloid-beta and tau (Ranasinghe et al., 2022). Studies in both humans and animal models support this finding of disrupted excitation-inhibition ratio as the disease progresses (Busche et al., 2012; Korzhova et al., 2021; van Nifterick et al., 2023; Zott et al., 2019). Moreover, disrupted intrinsic functional connectivity patterns of subregions of the hippocampus, nucleus accumbens, and amygdala are prominent in AD and linked to lower cognitive scores and synaptic dysfunction (Gonzalez-Rodriguez et al., 2023; Nie et al., 2017; Ortner et al., 2016; Poulin et al., 2011). Together, this body of work demonstrates that neural activity patterns are greatly altered across the progression of AD. However, how activation of brain regions and connectivity of neural networks might be influenced by parity with hAPOEε4 genotype at middle age remain unexplored, and thus, is the main question of the present study.

Building on previous work, the present study aims to address key gaps in the understanding about the combined effects of parity and hAPOEε4 genotype on the middle-aged brain. Our previous research identified unique signatures of primiparity and hAPOEε4, including increased use of a non-spatial cognitive strategy and decreased number and engagement of new-born neurons in the hippocampus (Lee et al., 2024). Despite these differences, overall performance in terms of errors committed in the spatial working memory task did not differ between groups at the end of testing. Importantly, this does not negate the possibility that we may see differences in underlying functional connectivity in wildtype and hAPOEε4 knock-in middle-aged female rats, and with previous parity. Here, this study seeks to elucidate how network-level dynamics contribute to cognitive outcomes and provide a more comprehensive view of neural mechanisms altered by primiparity at middle age in a model of AD risk.

## 2. Methods

### 2.1. Subjects

Forty-eight age-matched wildtype and humanized (h) APOEε4 knock-in (HsdSage:SD- ApoEem1Sage rat, developed by SAGE Labs, Inc., Saint Louis, MO, USA) Sprague Dawley female rats were used, as described in Lee et al. (2024). Briefly, rats were either nulliparous, with no sexual or reproductive experience and no exposure to pups, or primiparous, with one reproductive experience at 4 months of age. From pregnancy until postnatal day 45 in primiparous rats, all rats were single-housed, with primiparous and nulliparous rats in separate colony rooms to minimize any disruptions and cues from pups to nulliparous rats. Primiparous rats were housed with pups until weaning. Aside from this period, all rats were pair-housed. Rats were left undisturbed until middle age (12-13 months of age). All experiments were conducted in accordance with ethical guidelines set by the Canada Council for Animal Care, and all procedures were approved by the University of British Columbia Animal Care Committee. All efforts were made to reduce the number and suffering of animals.

### 2.2. Procedure

Starting at middle age (approximately 12-13 months old), rats were trained and tested using the delayed win-shift radial arm maze. For a detailed description of the cognitive task, refer to Lee et al. (2024). The task consisted of daily trials across 21 days, each comprising a training phase and a testing phase. During the training phase, 4 of the maze arms were randomly selected and baited with a quarter of a Froot Loop©, whereas the other 4 arms were blocked. Rats were allowed to explore the maze until all food rewards were retrieved or 5 minutes has elapsed. After the training phase, rats were returned to their home cages for a 5-minute delay (for the probe trial on the last day, day 21, the delay was increased to 30-minutes). In the subsequent testing phase, all maze arms were opened, but only the 4 arms that were previously blocked during training were baited with quarter of a Froot Loop©. All cognitive sessions were conducted between 11:00 h and 16:00 h. Performance on this task relies on the hippocampus and frontal cortex (Floresco et al., 1997), which are among the earliest regions to be affected in AD (DeTure & Dickson, 2019). As reported in Lee et al. (2024), hAPOEε4 rats committed more across-phase and within-phase errors in the cognitive task compared to wildtype rats (Supplemental Figure A2).

Starting at middle age, rats were vaginally lavaged daily until euthanasia. In both wildtype and hAPOEε4 rats, primiparous rats were less likely than nulliparous rats to display irregular estrous cycling, which was defined as consecutive lavage cycles varying in length and/or order of stages (Lee et al., 2024).

### 2.3. Tissue collection and processing

Ninety minutes after the last day of cognitive testing, rats were euthanized via lethal overdose of sodium pentobarbital. Brains were extracted and cut longitudinally into halves. The right hemispheres were flash frozen on dry ice and stored at −80 °C, and the left hemispheres were post-fixed for 24h in 4% paraformaldehyde (at 4 °C), then transferred to a 30 % sucrose solution for cyroprotection until the brains sank. Using a freezing microtome (2M2000R; Leica, Richmond Hill, ON), the left hemisphere of each brain was sliced into 35 μm coronal sections and collected in series of 5 throughout the frontal cortex and in series of 10 throughout the entire rostral-caudal extent of the hippocampus. Sections were stored in a cryoprotective medium (consisting of 0.1 M PBS, 30 % ethylene glycol, and 20 % glycerol) at −20 °C. Tissue were used for various immunohistochemistry and electrochemiluminescence assays, and this data is reported in (Lee et al., 2024).

For the present study, one series of frontal cortex sections and one series of hippocampus sections were stained for zinc finger-containing transcription factor 268 (zif268), which is an immediate early gene (IEG) critical for long-term potentiation and memory consolidation (Bozon et al., 2003; Jones et al., 2001). Free-floating tissue was thoroughly rinsed (3 x 10 minutes) in 0.1M PBS (pH 7.4) before staining and between each of the following procedures. Tissue was first blocked with blocking solution containing 3% normal donkey serum and 0.3% Triton-X in 0.1M PBS for 30 minutes. Tissue was incubated in primary antibody solution containing 1:250 rabbit anti-zif268 (Egr-1 SC-189; Santa Cruz Biotechnology, Santa Cruz, CA, USA) in a dilution solution containing 1% normal donkey serum and 0.3% Triton-X in 0.1M PBS for 48 hours at 4°C. Next, tissue was blocked with blocking solution for 30 minutes, then incubated in secondary antibody solution containing 1:500 donkey anti-rabbit Alexa Fluor 594 (Vector Laboratories) in a dilution solution containing 1% normal donkey serum and 0.3% Triton-X in 0.1M PBS. Lastly, tissue was incubated in 1:10000 DAPI for 2.5 minutes, then mounted onto slides and cover-slipped with PVA DABCO.

One series of hippocampal sections was double stained for DCX and zinc finger-containing transcription factor 268 (zif268), an immediate early gene (IEG) required for long-term potentiation and memory consolidation (Bozon et al., 2003). Tissue was thoroughly rinsed (3 x 10 minutes) in 0.1M PBS (pH 7.4) before staining and between each of the following procedures. Tissue was first blocked with blocking solution containing 3% normal donkey serum and 0.3% Triton-X in 0.1M PBS for 30 minutes. Tissue was incubated in primary antibody solution containing 1:500 goat anti-DCX (Santa Cruz Biotechnology, Santa Cruz, CA, USA) and 1:1000 rabbit anti-zif268 (Egr-1 SC-189; Santa Cruz Biotechnology, Santa Cruz, CA, USA) in a dilution solution containing 1% normal donkey serum and 0.3% Triton-X in 0.1M PBS for 24 hours at 4°C. Then, tissue was blocked with blocking solution for 30 minutes, then incubated in secondary antibody solution containing 1:500 donkey anti-goat Alexa Fluor 488 (Invitrogen) and 1:500 donkey anti-rabbit Alexa Fluor 594 (Vector Laboratories) in a dilution solution containing 1% normal donkey serum and 0.3% Triton-X in 0.1M PBS. Lastly, sections were mounted onto slides and cover-slipped with PVA DABCO.

### 2.4. Microscopy and cell quantification

Microscopy and cell quantification was conducted by an investigator blinded to experimental conditions of animals. Sections stained with zif268 were imaged using the Zeiss Axio Scan 7 (Carl Zeiss Microscopy, Thornwood, NY, USA) with a 20x objective lens using fluorescent imaging. Digitized images were analyzed to measure zif268-immunoreactive (IR) cell density using a MATLAB (MathWorks) code originally developed by JEJS and later optimized by BHL and RK. This code can be made available by contacting the corresponding author. Briefly, after loading an image, the investigator is prompted to manually trace regions of interest in the image. For each region of interest, the background (i.e. the region outside the region of interest) is removed and the region of interest is processed and binarized. Size restrictions are applied to remove artifacts that are too small or too large to be considered a zif268- IR cell. Within each region of interest, the number of cells is divided by the total area of the region of interest to calculate the cell density. Adjustments were made to the number of times the background was removed (between 2.75 and 8 times, depending on the section) and cell size restrictions (200 to 1000 pixels) to compensate for differences in staining intensity.

Two sections of each of the following brain regions of interest were quantified for zif268-IR cells: infralimbic cortex (IL), prelimbic cortex (PrL), anterior cingulate cortex (ACC), medial dorsal striatum (mDS), lateral dorsal striatum (lDS), nucleus accumbens core (NAc), nucleus accumbens shell (NAs), central nucleus of the amygdala (CeA), basolateral nucleus of the amygdala (BLA), lateral nucleus of the amygdala (LA), dorsal (d) hippocampus (dCA1, dCA3, dDG), ventral (v) hippocampus (vCA1, vCA3, vDG), granular retrosplenial cortex (subdivided into RSGc and RSGab), and agranular retrosplenial cortex (RSA). Brain regions were defined according to a standard rat brain atlas (Paxinos & Watson, 2006). These regions were chosen because of their involvement with performance in the cognitive task and association with AD-related changes in excitability (Cordella et al., 2018; Dickerson et al., 2005; Floresco et al., 1997; Palop & Mucke, 2016).

### 2.5. Statistical Analyses

Analysis of variance (ANOVA) were conducted using Statistica (Statsoft Tulsa, OK) and were used on regional densities of zif268-IR cells as the dependent variables of interest, with genotype (wildtype, hAPOEε4) and parity (nulliparous, primiparous) as between-subjects variables. Repeated measures ANOVA were used with subregion (grouped as: dorsal hippocampus (dCA1, dCA3, dDG), ventral hippocampus (vCA1, vCA3, vDG), nucleus accumbens (NAc, NAs), frontal cortex (IL, PrL, ACC), amygdala (CeA, BLA, LA), dorsal striatum (mDS, lDS), and retrosplenial cortex (RSA, RSGc, RSGab)) as within-subjects variables. Post-hoc tests used Newman-Keuls and any a priori comparisons were subjected to Bonferroni correction. Significance level was set at α=0.05. Effect sizes were calculated as partial η^2^ or Cohen’s d where appropriate.

#### 2.5.1. Pairwise correlation analysis and network visualization

Data was loaded into R, from which brain region data was isolated from animals in each of the four groups. Pairwise correlation matrices using Pearson’s method were calculated considering pairwise complete observations to account for any missing data. Heatmaps of the correlation matrices were generated for each group using the ggplot2 and reshape2 packages with a minimum |r| > 0.5.

Networks for each group were built using the igraph package, where nodes represent brain regions and edges represent correlations strong correlations. Edge weights were included only if |r| > 0.5. Brain subregions were grouped together into their associated brain regions and colour coding to enhance the visual distinction in the resulting network plots (Table 1).

**Table 1.** Grouping of brain regions of interest.

| Brain Regions | Brain Subregions |
| --- | --- |
| Dorsal hippocampus | "dCA1", "dCA3", "dDG", "dDG DCX" |
| Ventral hippocampus | "vCA1", "vCA3", "vDG", "vDG DCX" |
| Nucleus accumbens | "NAc", "NAs" |
| Frontal cortex | "PrL", "IL", "ACC" |
| Amygdala | "CeA", "BLA", "LA" |
| Dorsal Striatum | "mDS", "IDS" |
| Retrosplenial cortex | "RSA", "RSGc", "RSGab" |

The networks were represented by both a circular format for interpretable visualization of brain region and subregion differences between groups, as well as through a force directed layout based on |r| > 0.5 equivalent edge values. The edge width is relative to the maximum weight in each plot to standardize the visual impact of different correlation strengths across each network.

#### 2.5.2. Network metric analysis

Beyond examination of neural activity changes, studies have demonstrated the utility of using graph analytical measures to reveal specific features that are altered in a functional brain network (Binnewijzend et al., 2014; Lorenzini et al., 2023). Connectivity parameters of neural networks can reveal important context about the efficiency and integration of communication between brain regions (Scharwächter et al., 2022). As such, application of graph theory to understand functional connectivity provides the added benefit of uncovering common organizational principles that might underlying neural activation patterns, which would greatly enhance interpretation of activation data from individual regions or between pairs of regions.

From the force directed network, various centrality and clustering measures were extracted. These measures include eigenvector centrality, which describe the importance and connectivity of nodes within the network, as well as path length, density, global clustering coefficient, and modularity, which are network-wide metrics. Analyses focused on eigenvector values of nodes, which consider both the level of connectivity a brain region has as well as the connectivity of connected regions, thus representing overall influence and importance. These metrics were visualized using ggplot in R for individual brain regions as well as averaged across regions, including an error bar of the standard deviation, which captures the heterogeneity of subregion values.

#### 2.5.3. Brain region-behaviour correlations and interactions

Data was imported into R with packages tidyverse, glmnet, and corrplot loaded for data management, analysis, and visualization. The data standardized and split into four groups based on parity and genotype. A systematic approach was implemented to explore all combinations of up to four brain regions as predictors of behaviour (total number of across-phase and within-phase errors made on the last day of the cognitive task) (Lee et al., 2024). Multiple linear regression modelling was handled in R using the lm function, identifying behavior to be modelled as a dependent variable influenced by brain regions for each group. Results were extracted following each run of individual correlation calculations including adjusted R^2^ which accounts for the number of model predictors and statistical significance.

Brain region measurements were standardized followed by linear regression modelling now with interaction terms, including the main effects of group, region, and interactions to identify where predictive power of brain regions differs across groups. After fitting the linear model, a type 1 two-way ANOVA was performed to test for statistically significant interactions between regions and group correlations with behavior. This was performed for both individual and pairs of regions. All statistical analyses were performed in R (version 4.4.2).

## 3. Results

### 3.1.1. Primiparous hAPOEε4 rats had the lowest density of zif268-IR cells in the dorsal striatum and nucleus accumbens compared to all other groups

Primiparous hAPOEε4 rats had lower density of zif268-IR cells compared to all other groups in the dorsal striatum (parity and genotype interaction: *F*(1,32)=3.986, *p*=0.050, partial η^2^=0.111; Figure 1A,B), as well as in the nucleus accumbens (parity and genotype interaction: *F*(1,32)=4.849, *p*=0.035, partial η^2^=0.132; Figure 1C,D). There was a significant main effect of genotype, with hAPOEε4 rats showing lower density of zif268-IR cells compared to wildtype rats, in the dorsal striatum (*F*(1,32)=6.443, *p*=0.016, partial η^2^=0.168; Figure 1A,B) and nucleus accumbens (*F*(1,32)=7.518, *p*=0.010, partial η^2^=0.190; Figure 1C,D). There was also a significant main effect of region within the nucleus accumbens, with lower density of zif268-IR cells in the core than the shell (*F*(1,32)=29.035, *p*<0.001, partial η^2^=0.476; Figure 1C,D). There were no other significant main or interaction effects in the dorsal striatum (all *p*’s>0.068) or nucleus accumbens (all *p*’s>0.117).

**Figure 1.**
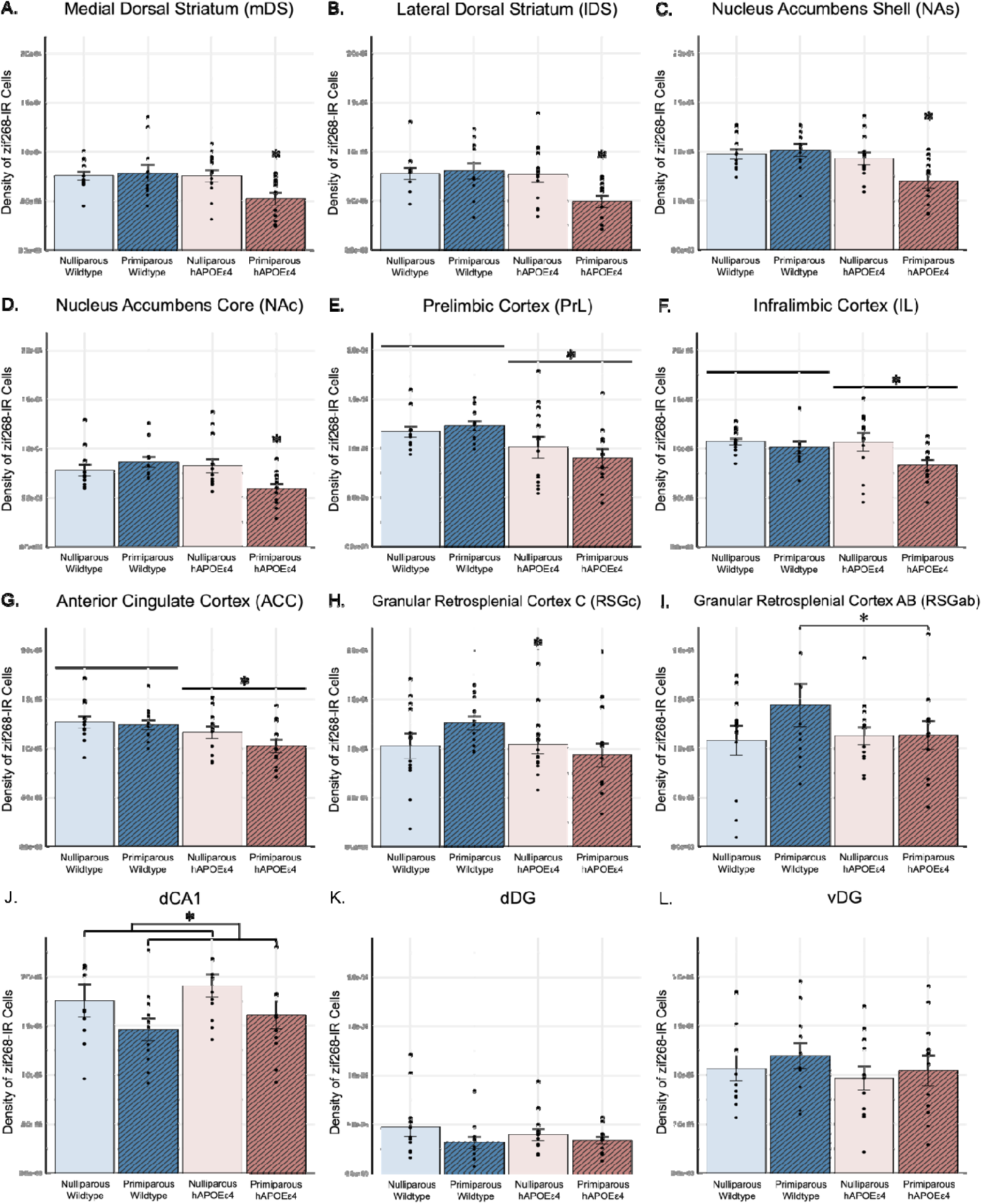
Average density of zif268-IR cells ± standard error of the mean in the (A) medial dorsal striatum, (B) lateral dorsal striatum, (C) nucleus accumbens shell, and (D) nucleus accumbens core. Zif268 cell density was the lowest in primiparous hAPOEε4 rats compared to all other groups in all subregions of the dorsal striatum and nucleus accumbens. Average density of zif268-IR cells ± standard error of the mean in the (E) prelimbic cortex, (F) infralimbic cortex, and (G) anterior cingulate cortex. Across all subregions of the frontal cortex, zif268 cell density was lower in primiparous hAPOEε4 rats compared to primiparous wildtype rats and in hAPOEε4 rats compared to wildtype rats. Zif268 cell density was greater in the ACC compared to PrL and IL. Average density of zif268-IR cells ± standard error of the mean in the (H) granular retrosplenial cortex C and (I) granular retrosplenial cortex AB. Zif268 cell density was the lower in primiparous hAPOEε4 rats compared to primiparous wildtype rats in the granular retrosplenial cortex. Average density of zif268-IR cells ± standard error of the mean in the (J) dorsal CA1, (K) dorsal DG, and (L) ventral DG. Zif268 cell density in the dorsal CA1 was lower in primiparous rats compared to nulliparous rats. Of all hippocampal subregions, zif2668 cell density was lowest in the dentate gyrus, with lower density in the dDG compared to vDG. * indicates *p*<0.05. CA – cornu ammonis, DG – dentate gyrus, zif268 – zinc finger-containing transcription factor 268, IR – immunoreactive, hAPOEε4 – humanized APOEε4

### 3.1.2. Primiparous hAPOEε4 rats had lower density of zif268-IR cells in the frontal cortex and retrosplenial cortex compared to primiparous wildtype rats

Across the frontal cortex, hAPOEε4 rats had lower density of zif268-IR cells than wildtype rats (main effect of genotype: *F*(1,37)=8.115, *p*=0.007, partial η^2^=0.180), although this effect is strongest in the PrL (Figure 1E) and ACC (Figure 1G). Planned comparisons revealed that primiparous hAPOEε4 rats had lower density of zif268-IR cells compared to primiparous wildtype rats across all subregions of the frontal cortex (Cohen’s *d*=1.634, *p*=0.007; parity and genotype interaction: *F*(1,37)=1.651, *p*=0.207, partial η^2^=0.043; Figure 1E,F,G). Density of zif268-IR cells was greater in the ACC compared to PrL and IL (main effect of region: *F*(2,74)=6.039, *p*=0.004, partial η^2^=0.140; Figure 1E,F,G). There were no other significant main or interaction effects on density of zif268-IR cells in frontal cortex subregions (all *p*’s>0.133).

Across the retrosplenial cortex, planned comparisons revealed that primiparous hAPOEε4 rats had lower density of zif268-IR cells compared to primiparous wildtype rats in the RSGc (Cohen’s *d*=0.951, *p*=0.014, Figure 1H) and RSGab (Cohen’s *d*=0.549, *p*=0.014, Figure 1I; parity by genotype by region interaction: *F*(1,76)=0.851, *p*=0.431, partial η^2^=0.022). There were no other significant main or interaction effects (all *p*’s>0.090).

### 3.1.3. Primiparous rats had lower density of zif268-IR cells in the dorsal CA1 compared to nulliparous rats

Primiparous rats had lower density of zif268-IR cells in the dorsal CA1 compared to nulliparous rats (parity and region interaction: *F*(5,180)=2.821, *p*=0.018, partial η^2^=0.073; Figure 1J). The density of zif268-IR cells was lowest in the dDG and vDG compared to all other subregions, with lower density in the dDG compared to vDG (main effect of region: *F*(5,180)=67.903, *p*=0<0.001, partial η^2^=0.654; Figure 1K,L). There were no other significant main or interaction effects on density of zif268-IR cells in hippocampal subregions (all *p*’s>0.667).

### 3.2. Functional Connectivity

#### 3.2.1. Correlation patterns of zif268 expression across brain regions of interest varied by primiparity and hAPOEε4 genotype

Pearson product-moment correlations were calculated with the densities of zif268-IR cells in 19 brain regions, as well as percentage of zif268/DCX co-labelled cells in the ventral and dorsal dentate gyrus to produce unique correlation matrices (where correlation coefficient |*r*| > 0.5) for each group (Figure 2A-D). Circular network maps were created as an alternative visualization of these correlations (Figure 2E-H).

**Figure 2.**
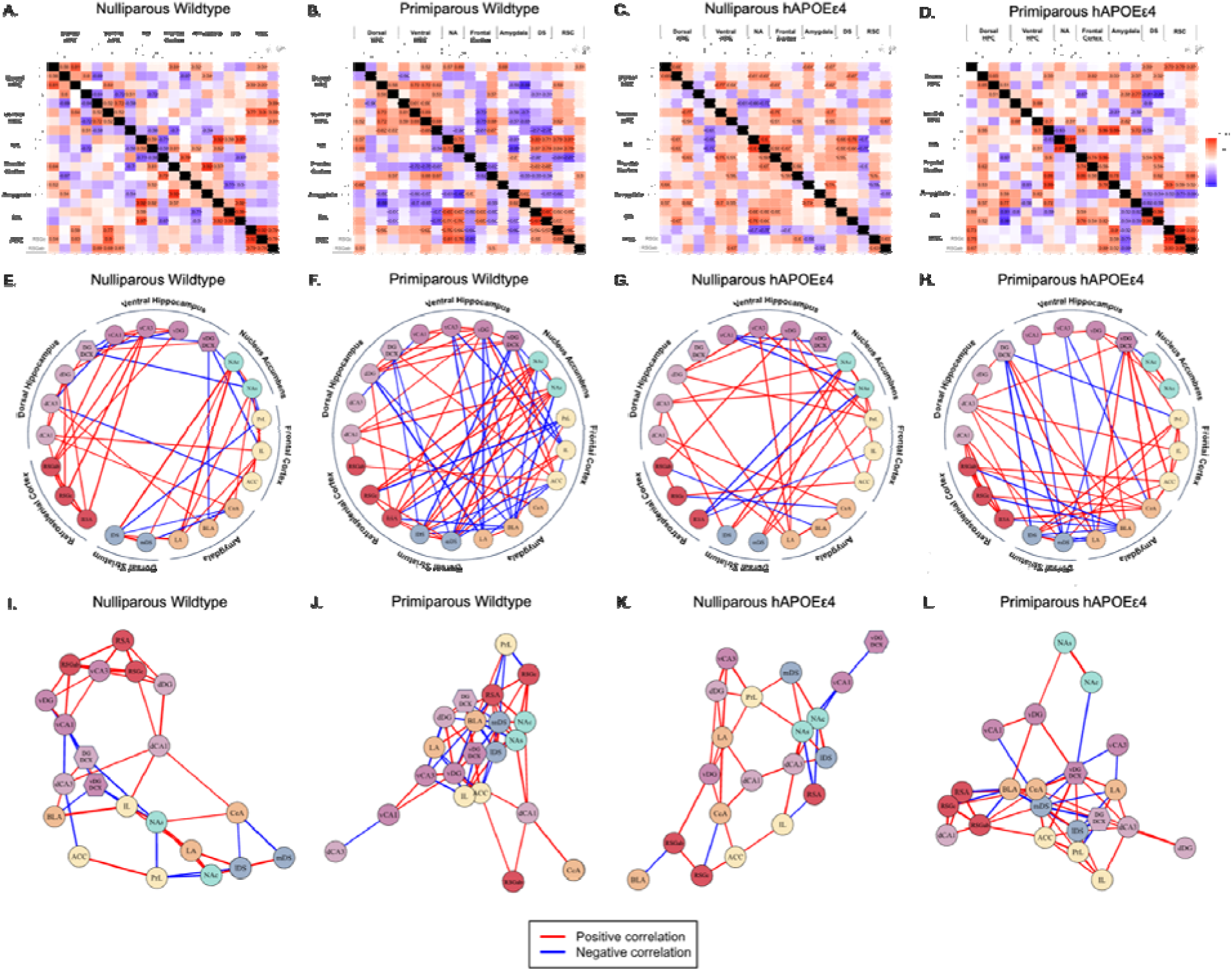
(A-D) Heat maps generated based on correlation coefficients between density of zif268-IR cells in the different regions of interest and percentage of DCX/zif268 co-labelled cells in the dorsal and ventral dentate gyrus across groups. All negative (blue) and positive (red) correlations are included, and boxes with text indicate those with a coefficient |*r*| > 0.5. (E-H) Circular network maps and (I-L) geometric network maps were generated based on correlation coefficients between density of zif268-IR cells in the different regions of interest and percentage of DCX/zif268 co-labelled cells in the dorsal and ventral dentate gyrus across groups. Negative (blue) and positive (red) correlations with coefficient |*r*| > 0.5 were included. hAPOEε4 – humanized APOEε4

#### 3.2.2. Connectivity networks of wildtype primiparous rats had the shortest path lengths, greatest density, highest clustering, and lowest modularity compared to all other groups, indicating greater efficacy and connectivity. hAPOEε4 genotype reduced clustering and network density, particularly in the network of primiparous rats

A network analysis revealed unique connectivity networks between regions within each group (Figure 2I-L). Primiparous wildtype rats had the shortest average path length between all brain regions, indicating greater efficiency within the network (Table 2). Primiparous wildtype rats also had the greatest overall network density, meaning a higher proportion of connections relative to the maximum possible connections (Table 2). hAPOEε4 genotype reduced network density in both nulliparous and primiparous rats (Table 2). Global clustering coefficients, which are based on the number of times three regions are communicating (A -> B -> C -> A), show that networks in primiparous wildtype rats were more highly clustered than networks in nulliparous wildtype rats, and hAPOEε4 genotype reduced the clustering effect, particularly in primiparous rats (Table 2). The modularity parameter, which determines the degree to which the network is segregated into non-overlapping modules or “subnetworks”, shows that region connectivity was more modular in networks of nulliparous rats than in networks of primiparous rats (Table 2).

**Table 2.** Network wide analysis metrics, including average path length, density, global clustering coefficient, and modularity. hAPOEε4 – humanized APOEε4.

| Group | Average Path Length | Density | Global Clustering | Modularity |
| --- | --- | --- | --- | --- |
| **Nulliparous Wildtype** | 1.436637742 | 0.228571429 | 0.407608696 | 0.444066136 |
| **Primiparous Wildtype** | 1.183815341 | 0.304761905 | 0.49009901 | 0.289302147 |
| **Nulliparous hAPOEε4** | 1.563981275 | 0.189473684 | 0.355932203 | 0.414589734 |
| **Primiparous hAPOEε4** | 1.264392998 | 0.252380952 | 0.382239382 | 0.332447347 |

#### 3.2.3. Brain regions were differentially important, and with varying heterogeneity across subregions, to connectivity networks depending on primiparity and hAPOEε4 genotype

Eigenvector centrality values (EV) were calculated for each region in each network based on the number of connections a region has, as well as the number of connections its neighbours have. As such, a high EV is representative of high importance of the given region within its network. Subregions were grouped together (for example, dCA1, dCA3, dDG, and dDG DCX were grouped together as part of the dorsal hippocampus), and the average EV for each group of subregions and standard deviations were calculated to represent the heterogeneity of region importance within each network.

In nulliparous wildtype rats, the regions with the highest average EVs were the retrosplenial cortex, followed by the dorsal and ventral hippocampus (Figures 3A-C,H). In nulliparous hAPOEε4 rats these regions had comparatively lower EVs, and the nucleus accumbens had the highest EV (Figure 3A,D). Primiparous rats had higher average EV for the frontal cortex and dorsal striatum compared to nulliparous rats, regardless of genotype (Figures 3A,E,G). The nucleus accumbens had a high average EV in primiparous wildtype rats yet a low average EV in primiparous hAPOEε4 rats (Figure 3A,D). Primiparous hAPOEε4 rats had higher average EVs for the frontal cortex and amygdala compared to all other groups (Figure 3A,E,F).

**Figure 3.**
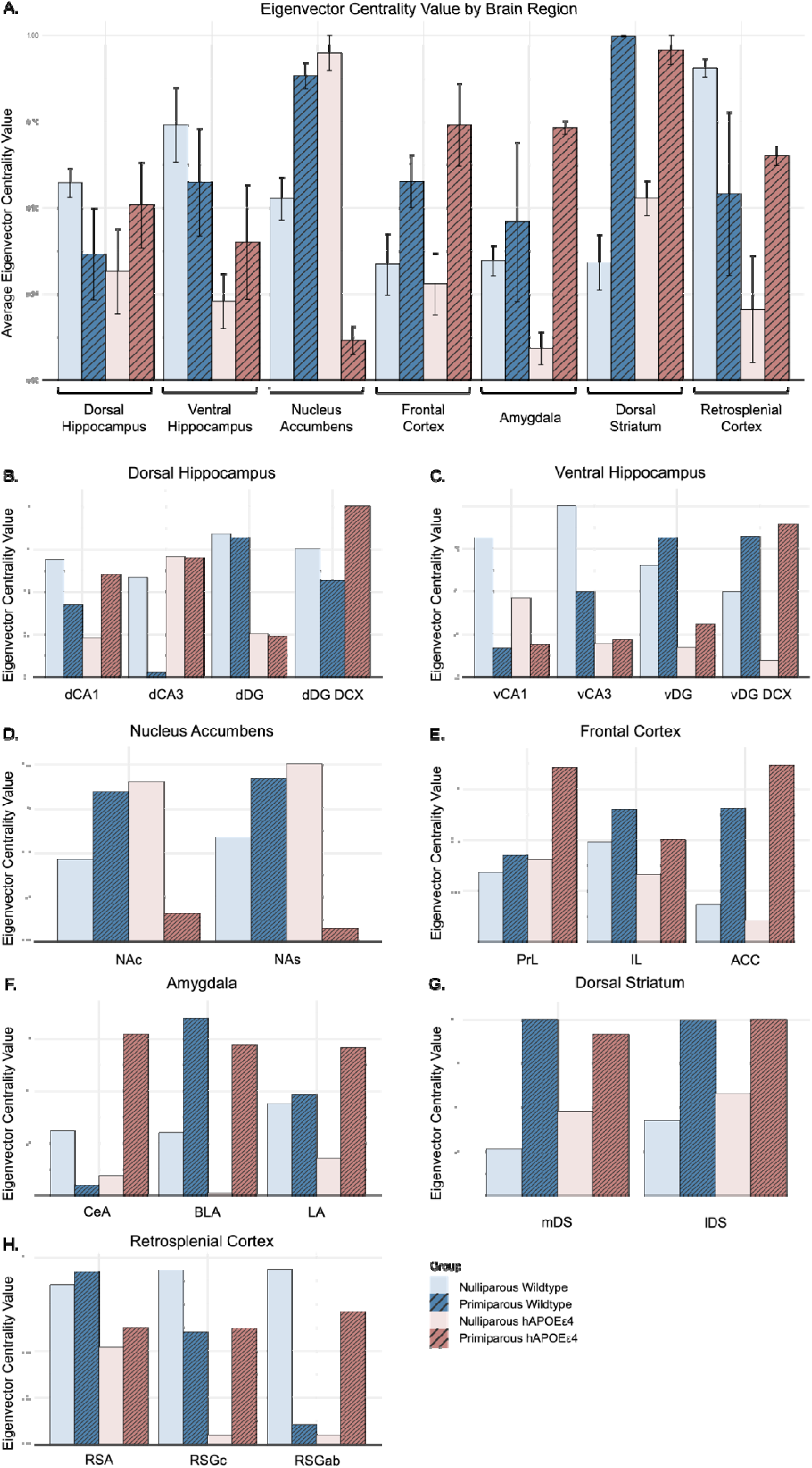
Eigenvector centrality values of the different subregions. A region with a high eigenvector centrality value indicates that region is highly important within its network. hAPOEε4 – humanized APOEε4

Next, EV standard deviations and EV of individual subregions were assessed to understand how importance of a region might be differentially distributed across its subregions depending on primiparity and hAPOEε4 genotype. In the nucleus accumbens, dorsal striatum, and frontal cortex, all groups showed similar EV standard deviations, indicating stable homogeneity of subregion importance across groups (Figure 3A). On the other hand, there was greater heterogeneity in subregion importance across groups in the dorsal and ventral hippocampus, retrosplenial cortex, and amygdala as seen in Figure 3A and described in the supplementary material.

### 3.3. Behaviour prediction of functional connectivity

To investigate the relationship between activation of regions of interest and behaviour, we employed a systematic analysis of regional connectivity across all brain regions of interest. The number of total errors made on the last day of the behavioural paradigm was used as the outcome variable, referred to as “errors” hereafter (see Lee et al. (2024) and the supplementary material for details). Importantly, there were no significant main or interaction effects on total errors made on the last day of the behavioural paradigm (all *p*’s>0.280; Figure 4). We systematically evaluated all combinations of up to four brain regions as predictors of errors using multiple linear regression (see supplementary material for details).

**Figure 4.**
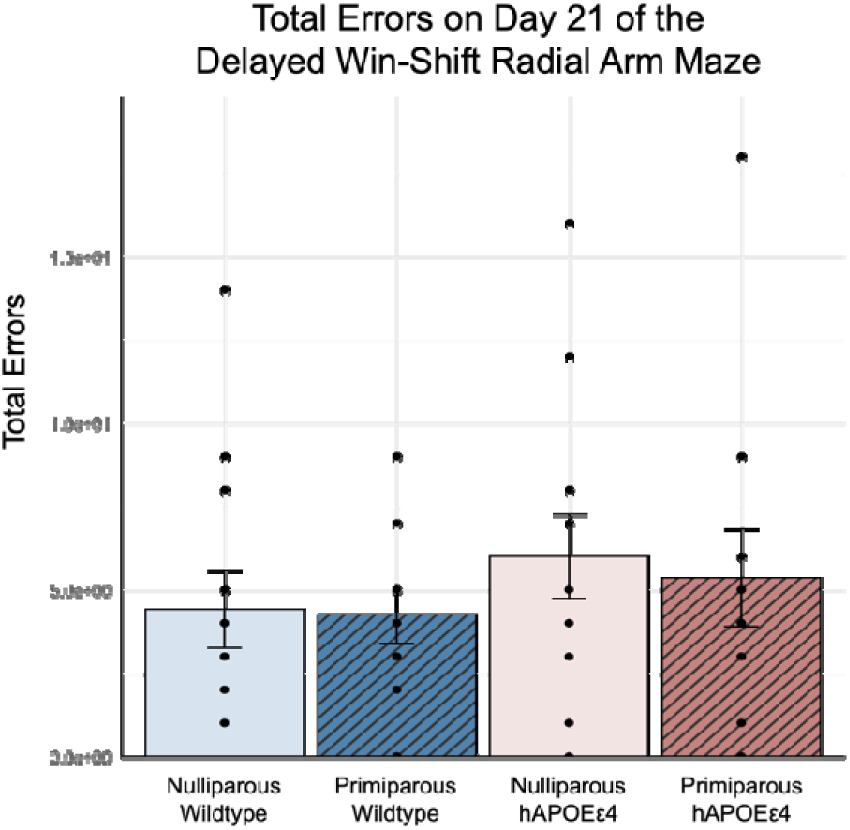
Total errors on day 21 of the behavioural paradigm. hAPOEε4 – humanized APOEε4

#### 3.3.1. In nulliparous wildtype rats, activation in mDS was negatively correlated with errors, whereas in primiparous wildtype rats, activation in mDS and NAc were positively correlated with errors. In nulliparous hAPOEε4 rats, activation in ACC was negatively correlated with errors. In primiparous hAPOEε4 rats, activation in vCA3 was negatively correlated with errors and activation in CeA was positively correlated with errors

Analysis of single region correlations with errors in each group revealed group-specific regional outcomes. In nulliparous wildtype rats, activation in the mDS was negatively correlated with errors (r=- 0.593, *p*=0.04; Figure 5), whereas in primiparous wildtype rats, activation in the mDS (R=0.697, *p*=0.017; Figure 5) and NAc (r=0.661, *p*=0.027; Figure 5) were positively correlated with errors. Activation in the lDS, LA, RSA, RSGc, and RSGab were also predictive of errors in nulliparous and primiparous wildtype rats in opposing directions, although these correlations were not significant (all p’s>0.085). In nulliparous hAPOEε4 rats, activation in the ACC was negatively correlated with errors (r=-0.610, *p*=0.027; Figure 5). Lastly, in primiparous hAPOEε4 rats, activation in the vCA3 was negatively correlated with errors (r=-0.646, *p*=0.032; Figure 5), and activation in the CeA was positively correlated with errors (r=0.691, *p*=0.019; Figure 5).

**Figure 5.**
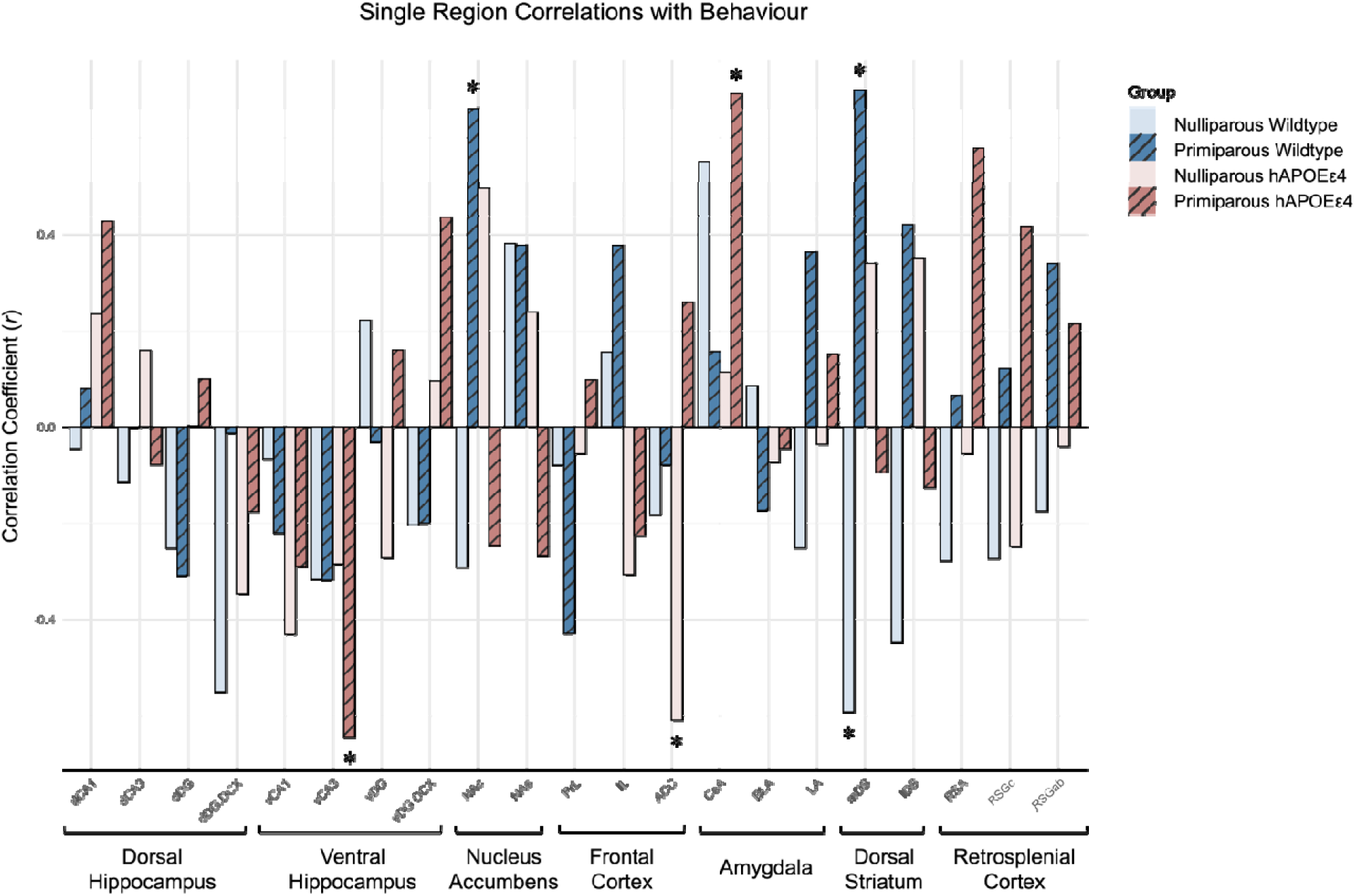
Single region correlations with behaviour, which was operationalized as the total number of errors made on day 21 of the behavioural paradigm. * indicates *p*<0.05. CA – cornu ammonis, DG – dentate gyrus, DCX – doublecortin, IR – immunoreactive, NAc – nucleus accumbens core, NAs – nucleus accumbens shell, PrL – prelimbic cortex, IL – infralimbic cortex, ACC – anterior cingulate cortex, CeA – central amygdala, BLA – basolateral amygdala, LA – lateral amygdala, mDS – medial dorsal striatum, lDS – lateral dorsal striatum, RSA – agranular retrosplenial cortex, RSG – granular retrosplenial cortex C (RSGc), RSGab – granular retrosplenial cortex AB, hAPOEε4 – humanized APOEε4

#### 3.3.2. Activation in region pairs involving subregions of the dorsal striatum and hippocampus was negatively correlated with errors in nulliparous wildtype rats and positively correlated with errors in primiparous wildtype rats. In nulliparous hAPOEε4 rats, activation in region pairs involving the ACC was negatively correlated with errors, while activation in NAc and RSA was positively correlated with errors. In primiparous hAPOEε4 rats, activation in vDG DCX and RSA was positively correlated with errors and activation in vCA3 and NAc was negatively correlated with errors

To examine group differences in how a region correlated with errors, linear models were produced to test for statistically significant interactions (parity by genotype). No brain regions individually showed significant interactions between parity and genotype (all *p*’s>0.065).

Activation in region pairs involving the dorsal striatum and DG new-born neurons or ventral CA3 correlated with errors in wildtype rats only, and the direction of the correlations depended on parity (interactions between parity and genotype: mDS-dDG DCX (F=3.327, *p*=0.029; Figure 6A), mDS-vCA3 (F=2.944, *p*=0.045; Figure 6C), mDS-vDG DCX (F=3.501, *p*=0.024; Figure 6D), and lDS-dDG DCX ((F=3.052, *p*=0.0397; Figure 6B). Activation in these region pairs involving dorsal striatum and DG new-born neurons were negatively correlated with errors in nulliparous wildtype rats (mDS and dDG DCX: *r*=- 0.703, *p*=0.011; lDS and dDG DCX: *r*=-0.660, *p*=0.019; mDS and vCA3: *r*=-0.641, *p*=0.025) but positively correlated with errors in primiparous wildtype rats (mDS and dDG DCX: *r*=0.630, *p*=0.038; mDS and vDG DCX: *r*=0.620, *p*=0.024; Figure 6D).

**Figure 6.**
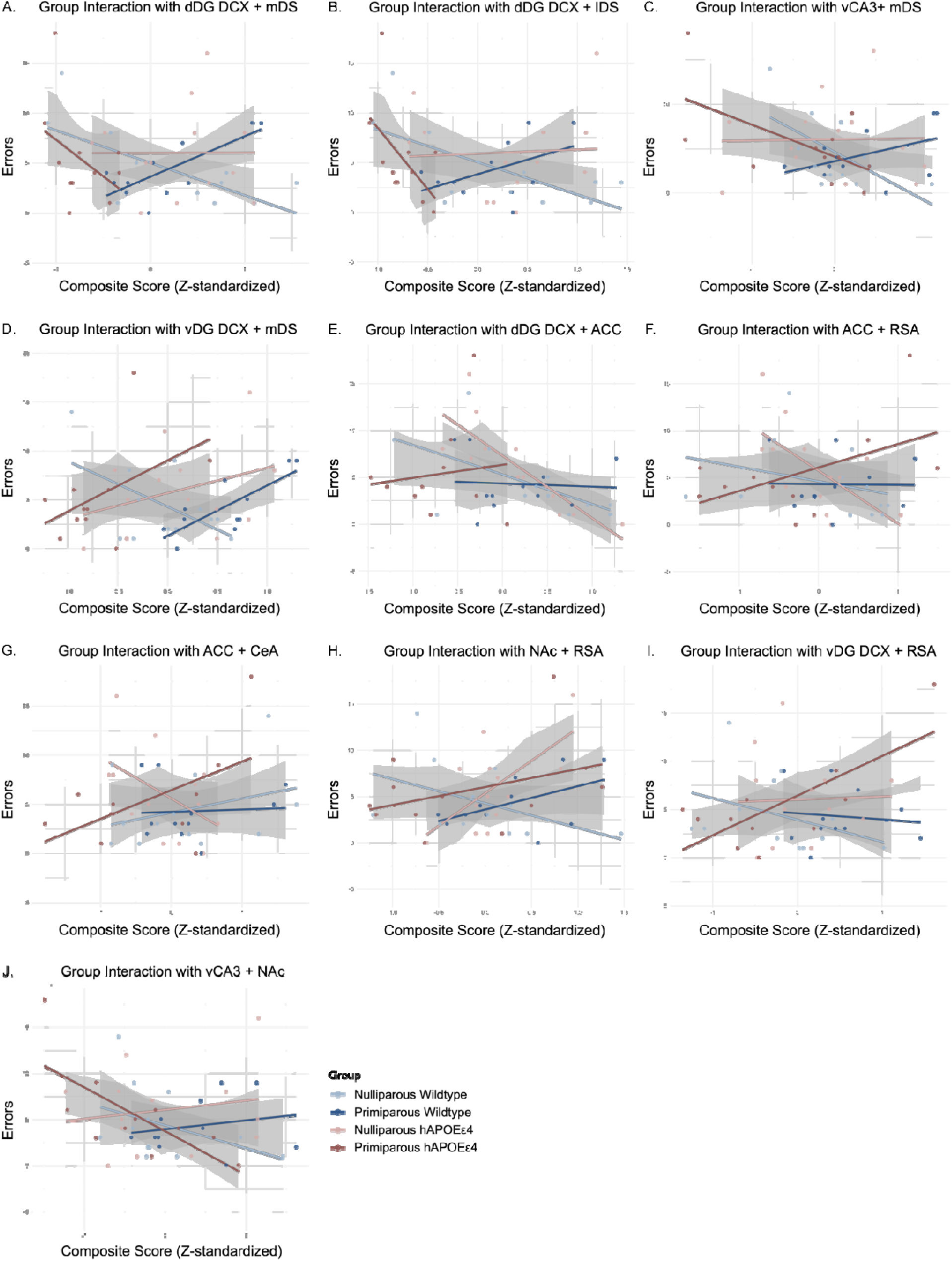
Linear models of interactions between behaviour, operationalized as the total number of errors made on day 21 of the behavioural paradigm, and (A) mDS and dDG DCX, (B) lDS and dDG DCX, (C) mDS and vCA3, (D) mDS and vDG DCX, (E) ACC and dDG DCX, (F) ACC and RSA, (G) ACC and CeA, (H) NAc and RSA, (I) vDG DCX and RSA, and (J) vCA3 and NAc. CA – cornu ammonis, DG – dentate gyrus, DCX – doublecortin, NAc – nucleus accumbens core, ACC – anterior cingulate cortex, CeA – central amygdala, mDS – medial dorsal striatum, lDS – lateral dorsal striatum, RSA – agranular retrosplenial cortex, hAPOEε4 – humanized APOEε4

Activation in region pairs that correlated with behaviour and differed in nulliparous hAPOEε4 rats involved the ACC and dDG new-born neurons, retrosplenial cortex, or central amygdala. Activation in region pairs involving the ACC was negatively correlated with errors in nulliparous hAPOEε4 rats (ACC and dDG DCX: *r*=-0.913, *p*=0.001; ACC and RSA: *r*=-0.567, *p*=0.043; interactions between parity and genotype: ACC-dDG DCX (F=2.900, *p*=0.0470; Figure 6E), ACC-RSA (F=4.190, *p*=0.012; Figure 6F), and ACC-CeA (F=2.857, *p*=0.049; Figure 6G). Activation in the region pair NAc-RSA was positively correlated with errors in nulliparous hAPOEε4 rats (*r*=0.563, *p*=0.045; interactions between group and genotype F=3.555, *p*=0.023; Figure 6H).

Region pairs that were significantly different in primiparous hAPOEε4 compared to all other groups involved the ventral hippocampus, with a positive correlation between vDG DCX-RSA activation and errors (*r*=0.736, *p*=0.010; interaction: F=2.918, *p*=0.046; Figure 6I), and a negative correlation between vCA3-NAc activation and errors (*r*=-0.672, *p*=0.024;interaction of group by genotype F=2.901, *p*=0.045; Figure 6J).

## 4. Discussion

Overall, we found that primiparous hAPOEε4 rats presented with widespread reductions in neural activation across subregions of the dorsal striatum, nucleus accumbens, frontal cortex, and retrosplenial cortex. We identified that each group, depending on parity and genotype, presented with unique functional connectivity. Primiparous wildtype rats had networks with greater network efficiency and cohesiveness compared to all other groups. hAPOEε4 genotype reduced clustering in the network of primiparous rats, suggesting detrimental effects to network integration. Depending on parity and hAPOEε4 genotype, different regions emerged as more influential, with high eigenvector centrality, within their networks. For example, the hippocampus and retrosplenial cortex were the most influential or important regions within the network of nulliparous wildtype rats, and eigenvector of these regions were greatly reduced with hAPOEε4 genotype. The frontal cortex and dorsal striatum were both highly influential within the network of primiparous rats, regardless of genotype. The nucleus accumbens was one of the most influential regions within the network of primiparous wildtype rats, yet the least influential within that of primiparous hAPOEε4 rats.

Although understanding the effects of parity and hAPOEε4 on activation patterns across the brain is valuable, it is also important to assess how changes in functional connectivity relate to behaviour. As such, we examined which individual regions and pairs of regions were predictive of errors committed on the last day of the spatial working memory task and how these relationships differed by parity and hAPOEε4 genotype. We found that activation in dorsal striatum and nucleus accumbens subregions were highly predictive of errors in wildtype rats – predicting reduced errors in nulliparous rats but predicting increased errors in the primiparous rats. In hAPOEε4 rats, activation in different regions were predictive of errors, indicating a shift in the importance of brain regions underlying spatial working memory. In nulliparous hAPOEε4 rats, activation in the ACC, both alone and in conjunction with activation of hippocampal new-born neurons, RSA, or the CeA, was associated with reduced errors. In primiparous hAPOEε4 rats, activation in the vCA3 was associated with reduced errors, but activation in the CeA was associated with increased errors. Intriguingly, activation of hippocampal new-born neurons in conjunction with activation in other regions was predictive of errors in directions that changed depending on parity and hAPOEε4 genotype. Activation of new neurons in conjunction with the dorsal striatum was associated with reduced errors in nulliparous wildtype rats, but increased errors in primiparous wildtype rats. Activation of new neurons in conjunction with the ACC was associated with reduced errors in nulliparous hAPOEε4 rats, but activation of new neurons in conjunction with the RSA was associated with increased errors in primiparous hAPOEε4 rats. In sum, the findings of this study highlight the dynamic interplay between previous parity and hAPOEε4 in shaping neural activation and functional connectivity at middle age and offer new insight into their potential contributions to cognitive function and the aging brain.

### 4.1. Primiparous hAPOEε4 rats had comparatively lower activation across a number of regions compared to all other groups

Primiparous hAPOEε4 rats had the lowest activation in subregions of both the dorsal striatum and nucleus accumbens compared to all other groups. The dorsal striatum and nucleus accumbens are both integral parts of the mesolimbic reward system, which is involved in cognition as well as other behaviours related to reward processing and incentive-based learning – all of which are impacted in AD (D’Amelio et al., 2018; Daniel & Pollmann, 2014; Perry & Kramer, 2015). Primiparous hAPOEε4 rats also had lower activation in subregions of the frontal cortex and granular retrosplenial cortex compared to primiparous wildtype rats. In humans with MCI, hypometabolism in the brain, particularly in the retrosplenial cortex, was predictive of eventual development of AD – with greater effect sizes in human females than males – suggesting the retrosplenial cortex might be an early site of dysfunction in those at risk of developing AD (Terstege, Galea, et al., 2024). In the present study, we found reduced activation in the retrosplenial cortex with hAPOEε4 genotype, but only in primiparous rats, which indicates that previous parity may exacerbate the lowered neural activity (or hypometabolism) seen with increased AD risk, only in rats with hAPOEε4 genotype. Interestingly, the reduced activation in primiparous hAPOEε4 rats was only evident in the granular subdivision of the retrosplenial cortex, an area that receives direct inputs from the dorsal hippocampus and is involved in processing spatial and contextual information (Tsai et al., 2022). Collectively, these data illustrate that activation across a wide variety of brain regions was uniquely impacted in primiparous rats by hAPOEε4 genotype. This extends our previous findings on the “parity paradox”, showing that brain activation differs by parity depending on hAPOEε4 genotype (Lee et al., 2024).

Across frontal cortex, dorsal striatum, and nucleus accumbens subregions, neural activation was reduced in hAPOEε4 rats compared to wildtype rats, suggesting that these regions might be particularly vulnerable to neuronal excitability changes related to hAPOEε4 genotype at middle age. This might seem inconsistent with literature showing neuronal hyperexcitability in studies using animal models of AD or in vitro studies (Ghatak et al., 2019; Palop & Mucke, 2016; Šišková et al., 2014). However, much of this work has been focused on examining electrophysiological properties of isolated neurons or cultures, and no studies to date have investigated neural activation patterns across the brain in vivo in a model of late-onset sporadic AD risk in females. The human literature is mixed, with some studies reporting increased activity (Bookheimer et al., 2000) and some reporting decreased activity (Dickerson et al., 2005; Terstege, Galea, et al., 2024) with AD or AD risk – although most were centered around activity of the hippocampus and related regions. Nonetheless, apathy and decreased pursuit of rewarding behaviours have often been observed in individuals with AD, and have been associated with atrophy in areas including the ACC and medial prefrontal cortex, as well as altered dopaminergic activity in the nucleus accumbens (Koch et al., 2014; Perry & Kramer, 2015; Rosen et al., 2005). Therefore, it may not be entirely surprising that hAPOEε4 genotype reduced activation of these regions in the present study. It would be interesting for future research to explore influences of hAPOEε4 on the dopaminergic system and other components of the reward system.

There were minimal changes in activation with primiparity and hAPOEε4 across subregions of the hippocampus and amygdala, except for dorsal CA1, which showed reduced activation in primiparous rats compared to nulliparous rats, regardless of genotype. There is evidence to suggest that changes to neural activation might manifest at different stages of AD, as activation was increased in individuals with mild cognitive impairment compared to both those who were cognitive intact and diagnosed with AD (Dickerson et al., 2005). Indeed, there is support in the literature for a shift from hyperactivation to hypoactivation in the hippocampus as the disease progresses (Celone et al., 2006). In accordance with this idea, it is possible that assessing earlier and later time points in hAPOEε4 rats might reveal dynamic differences in neural activation within the hippocampus, and potentially other brain regions.

### 4.2. Primiparous wildtype rats had a more efficient, integrated, and cohesive neural network, and primiparous hAPOEε4 rats showed reduced clustering within the network

Functional connectivity networks generated using correlations of zif268 cell density across all regions of interest varied by several parameters, depending on primiparity and hAPOEε4 genotype. Compared to all other groups, the network of primiparous wildtype rats had a shorter average path length and greater density, which is indicative of greater network efficiency. The network of primiparous wildtype rats also displayed higher clustering and less modularity, which is indicative of less network fragmentation or segregation, and more integration. This is in line with literature showing enhanced performance on hippocampus-based cognitive tasks with parity history in both animal models and humans (Barha et al., 2015; Cui et al., 2014; Galea et al., 2018; Orchard et al., 2020). Together, this suggests that the long-lasting neuroprotective effects of previous parity extends beyond hippocampal integrity and cognition, to neural network organization at middle age.

Intriguingly, females present with greater age-related changes in the organization and efficiency of specific functional networks, and one notable example is seen in less segregation within the default mode network, which has been implicated in AD, compared to males (Anticevic et al., 2012; Ballard et al., 2022; Greicius et al., 2004). Functional network segregation has been suggested to be linked to specialized information processing and efficiency, and thus the decrease in age-related network segregation might be underlying functional decline (Ballard et al., 2022). In the present study, we found that although primiparous wildtype rats showed decreased segregation in the neural network, they also show greater clustering and increased network efficiency. These parameters together are suggestive of resilience or a compensatory response within the neural network of primiparous wildtype rats at middle age. In sum, these findings suggest that previous parity increased efficiency, integration, and cohesiveness in the neural network, enhancing overall functional connectivity, in wildtype rats at middle age.

The present study further identified that hAPOEε4 genotype reduced network clustering, and this effect was particularly pronounced in primiparous rats. This finding is consistent with other studies that have reported reduced clustering in functional connectivity networks in humans with AD compared to healthy controls, although these studies did not analyze their findings with sex as a factor (Brier et al., 2014; Supekar et al., 2008). More research is needed to systematically investigate how age and AD-related changes in topological features of neural networks might vary by sex and examine how sex-specific factors might further contribute to these relationships.

### 4.3. Parity was differentially associated with changes in importance of the hippocampus and retrosplenial cortex within the neural network, depending on hAPOEε4 genotype

In the present study, the retrosplenial cortex, dorsal hippocampus, and ventral hippocampus were the most important regions within the network of nulliparous wildtype rats, and importance of these regions were considerably reduced in the network of nulliparous hAPOEε4 rats. This is consistent with research in humans showing functional disconnection of the hippocampus with other parts of the brain as AD progresses (Dautricourt et al., 2021; Rao et al., 2022). In the present study, we further show that importance of these regions was lower in the network of primiparous wildtype than nulliparous wildtype rats, and higher in that of primiparous hAPOEε4 than nulliparous hAPOEε4 rats, which suggests an interaction between previous parity and hAPOEε4 genotype to alter how the hippocampus and retrosplenial cortex are activated with other regions in the brain in response to memory retrieval.

#### 4.3.1. The frontal cortex and dorsal striatum showed high importance within the neural network of primiparous rats. The nucleus accumbens showed high importance within the neural network of primiparous wildtype rats, but low importance within the neural network of primiparous hAPOEε4 rats

In the present study, the frontal cortex and dorsal striatum were more important in the network of primiparous rats compared to nulliparous rats, regardless of hAPOEe4 genotype. Previous research have demonstrated long-term influences of parity on brain aging and grey matter volume in humans (Aleknaviciute et al., 2022; de Lange et al., 2019) and hippocampal neurogenesis and cognition in rodents (Barha et al., 2015; Eid et al., 2019). The findings of the present study additionally show that previous parity leads to a shift in the neural network towards greater importance of the frontal cortex and dorsal striatum within the neural network, that are apparent even at middle age.

Interestingly, although activation of the dorsal striatum was reduced in primiparous hAPOEe4 rats compared to all other groups, importance of the dorsal striatum within the neural network was preserved in this group. One interpretation of this is that connectivity of the dorsal striatum with other brain regions is resilient against the reductions in neural activity related to primiparity and hAPOEe4 genotype at middle age. On the other hand, both activation and importance of the nucleus accumbens within the neural network was low in primiparous hAPOEe4 rats. This finding suggests that the nucleus accumbens may be particularly vulnerable to the combined effects of primiparity and hAPOEε4 genotype. The nucleus accumbens is a critical hub for integrating reward-related and goal-directed behaviors, and as such, reduced functional integration with other regions could be linked to alterations in motivation or decision-making processes, which has been observed in models of AD (Cordella et al., 2018). Future studies should investigate whether these changes in the nucleus accumbens are associated with specific behavioural or cognitive deficits in primiparous hAPOEε4 rats.

### 4.4. Activation of hippocampal new-born neurons in conjunction with the dorsal striatum, anterior cingulate cortex, or retrosplenial cortex were dynamically predictive of errors depending on primiparity and hAPOEε4 genotype

The present study aimed to elucidate the relationship between functional connectivity and cognitive performance by examining how brain regions predicted the number of errors made on the last day of the behavioural paradigm, depending on primiparity and hAPOEε4 genotype. Activation in the dorsal striatum, both alone and in conjunction with hippocampal new-born neurons, predicted errors in opposite directions depending on parity. Activation in the dorsal striatum and new neurons in the dDG was predictive of reduced errors in nulliparous wildtype rats but increased errors in primiparous wildtype rats, illustrating a shift with parity. This aligns with research indicating that the dorsal striatum and hippocampus employ distinct and complementary principles to support spatial navigation and memory (Ferbinteanu, 2020; Geerts et al., 2020). The findings of the present study suggest that primiparity confers lasting changes to the functional connectivity between the dorsal striatum and the hippocampus underlying spatial working memory. It is essential to emphasize that these differences emerge despite similar performance on the last day of the behavioural paradigm.

In nulliparous hAPOEε4 rats, activation in the ACC, both alone and in conjunction with other regions, was predictive of reduced errors, whereas activation in the region pair NAc-RSA was predictive of increased errors. In nulliparous hAPOEε4 rats, activation of new neurons in the dDG with ACC was predictive of reduced errors, whereas in primiparous hAPOEε4 rats, activation of new neurons in the vDG with RSA was predictive of increased errors. Together, this data demonstrates that activation of hippocampal new-born neurons in conjunction with different brain regions was predictive of errors made in the spatial working memory task, in directions that depended on parity and hAPOEε4 genotype. Moreover, activation in the agranular retrosplenial cortex – in conjunction with the NAc in nulliparous rats, or with new-born neurons in the vDG in primiparous rats – was predictive of increased errors in the spatial working memory task specifically with hAPOEε4 genotype, as these relationships were not observed in wildtype rats. This is consistent with literature previously mentioned, showing that hypometabolism and dysregulated functional connectivity of the retrosplenial cortex predicted conversion from mild cognitive impairment to AD in humans and correlated with impaired cognitive performance in both humans and rodent models (Mayne et al., 2024; Terstege et al., 2022; Terstege, Galea, et al., 2024). In sum, these findings highlight the nuanced and region-specific effects of primiparity and hAPOEε4 genotype on neural network connectivity and their functional implications.

### 4.5. Conclusions

Our findings demonstrate that primiparity in wildtype rats was associated with enhanced neural network efficiency and integration. In contrast, primiparity in hAPOEε4 rats was linked to reduced neural activation of key brain regions critical for memory and reward processing, including the dorsal striatum, nucleus accumbens, frontal cortex, and retrosplenial cortex. Primiparity in hAPOEε4 rats was additionally associated with increased fragmentation of the overall neural network. Among 19 regions of interest, activation of subregions in the dorsal striatum, nucleus accumbens, retrosplenial cortex, and hippocampus consistently predicted performance in a spatial working memory task, with these effects differing depending on primiparity and hAPOEε4 genotype. Interestingly, activation of hippocampal new-born neurons, in conjunction with the dorsal striatum, ACC, or retrosplenial cortex, was dynamically predictive of errors depending on primiparity and hAPOEε4 genotype. By revealing distinct patterns of network organization and regional importance, this study provides new insight into the neural mechanisms underlying cognitive outcomes altered by primiparity and hAPOEε4 in middle-aged female rats. This work underscores the power of investigating sex-specific variables and genetic risk factors for AD together to understand changes in brain health and their implications for cognitive resilience or vulnerability in aging populations.

## Notes

### Competing Interest Statement

The authors have declared no competing interest.

