## Supplement for "Parity and APOEε4 genotype contribute distinct changes to functional connectivity across the middle-aged brain"

**Supplementary Material**

**A1. Density of zif268-IR cells**

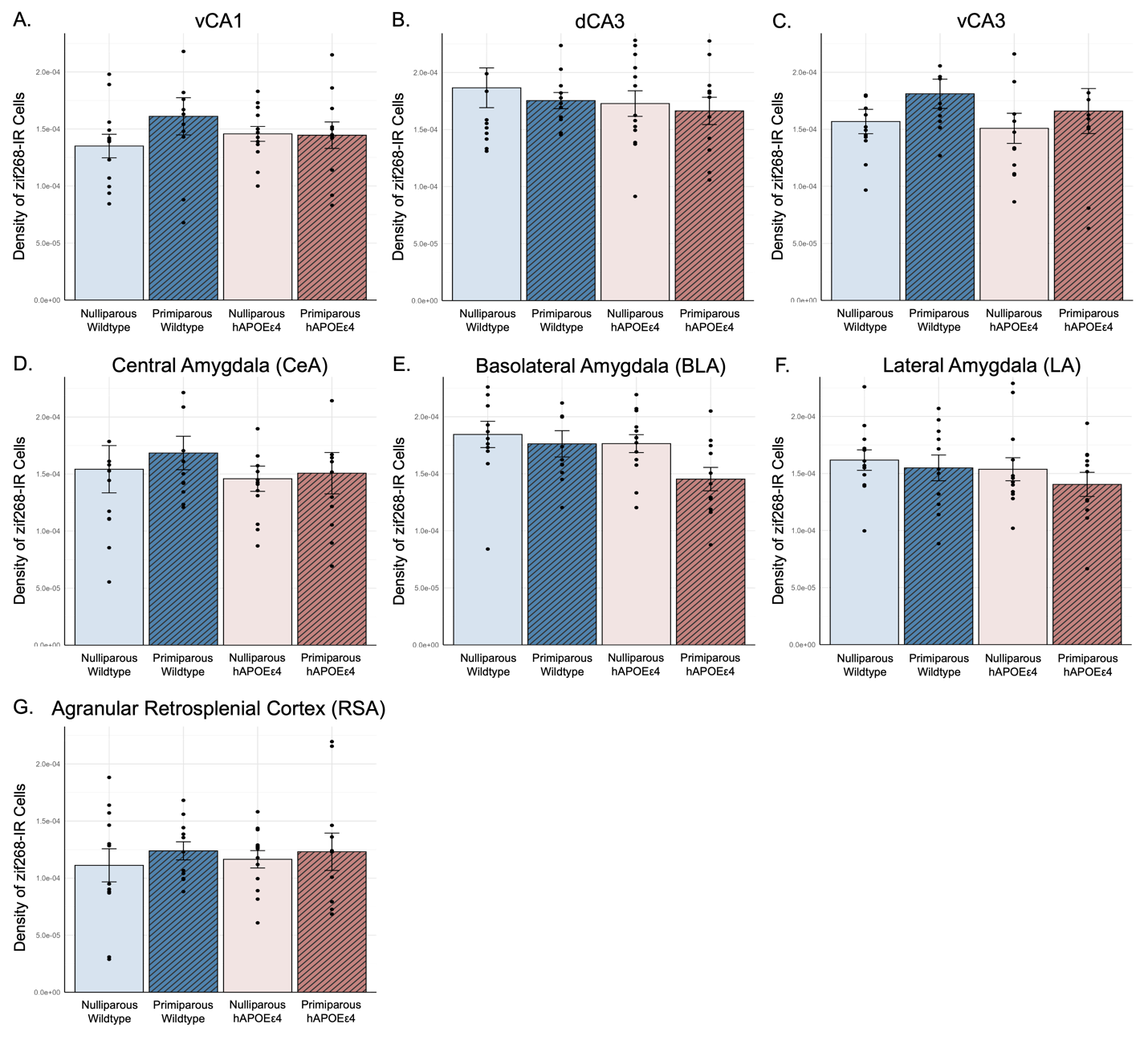

Figure A1. Average density of zif268-IR cells ± standard error of the mean in the (A) ventral CA1, (B) dorsal CA3, (C) ventral CA3, (D) central amygdala, (E) basolateral amygdala, (F) lateral amygdala, and (G) agranular retrosplenial cortex. zif268 – zinc finger-containing transcription factor 268, IR – immunoreactive, hAPOEε4 – humanized APOEε4

**A2. Eigenvector centrality for all regions of interest**

In the **dorsal hippocampus**, nulliparous wildtype rats showed the smallest EV standard deviation, as EVs across subregions of the dorsal hippocampus were stable across groups (Figure 3A). In wildtype rats, primiparity reduced the EV of dCA1 and dCA3, whereas in hAPOEε4 rats, primiparity increased the EV of dCA1 and the EV of dCA3 was increased regardless of primiparity (Figure 3B). The EV of dDG was reduced by hAPOEε4, with no notable differences by genotype between nulliparous and primiparous rats (Figure 3B). Relative to nulliparous wildtype rats, primiparity reduced the EV of dDG DCX cells and hAPOEε4 genotype increased it. dDG DCX is not present in the nulliparous APOEe4 network as it was not strongly correlated with any regions of interest (Figure 3B).

In the **ventral hippocampus**, hAPOEε4 rats, regardless of parity, had greater EV standard deviations compared to wildtype rats (Figure 3A). Primiparity greatly reduced the EV of vCA1, regardless of genotype, and hAPOEε4 reduced the EV of dCA1 only in nulliparous rats (Figure 3C). vCA3 was the most important node in the network of nulliparous wildtype rats (EV=1; Figure 3C). Primiparity reduced the EV in wildtype rats, and hAPOEε4 further reduced the EV of dCA3 in both nulliparous and primiparous rats (Figure 3C). Interestingly, the vDG and vDG DCX cells share a trend with higher EVs in primiparous rats compared to nulliparous rats (Figure 3C). Additionally, the EV of vDG was lower in hAPOEε4 rats compared to wildtype rats. Across the networks, vDG DCX had the lowest EV in nulliparous hAPOEε4 rats, but the highest EV in primiparous hAPOEε4 rats.

**Nucleus accumbens** subregions responded with a shared trend, whereby primiparity or hAPOEε4 resulted in increased EV compared to nulliparous wildtype rats (Figure 3D). On the other hand, in primiparous hAPOEε4 rats, the nucleus accumbens was least important in the network, with a notably lower EV, compared to all other groups (Figure 3D).

Within the **frontal cortex**, primiparity increased EVs – although to differing degrees across subregions (Figure 3E). Primiparity resulted in a marginal increase in PrL EV in wildtype rats, and a much larger increase in hAPOEε4 rats (Figure 3E). Primiparity increased IL EV in both wildtype and hAPOEε4 rats, although hAPOEε4 rats showed generally lower IL EVs compared to wildtype rats (Figure 3E). The ACC displayed drastic changes in EV with respect to primiparity and hAPOEε4 genotype, as the EV was low in nulliparous wildtype animals and even lower in nulliparous hAPOEε4 rats, and primiparity increased the EV in wildtype rats and increased the EV to an even greater extent in hAPOEε4 rats (Figure 3E).

In the **amygdala**, primiparous wildtype rats had the greatest EV standard deviation, with EV of the CeA being lowest and BLA being highest (Figure 3E), compared to relatively similar EV standard deviations in all other groups (Figure 3A). Compared to nulliparous wildtype rats, the CeA EV was reduced in primiparous wildtype rats and nulliparous hAPOEε4 rats (Figure 3F). However, CeA EV was greatly increased in primiparous hAPOEε4 rats (Figure 3F). The BLA EVs were lower in nulliparous rats compared to primiparous rats. In fact, the BLA in nulliparous hAPOEε4 rats was nearly completely disconnected from its network, with an EV of 0.009 (Figure 3F). BLA EV was high in primiparous rats, regardless of genotype (Figure 3F). The LA shared a similar pattern, although the differences were less pronounced: EV was reduced in nulliparous hAPOEε4 relative to nulliparous wildtype rats, and there was a marginal increase in LA EV with primiparity in wildtype rats and a major increase with primiparity in hAPOEε4 rats (Figure 3F).

hAPOEε4 genotype increased EVs of **dorsal striatum** subregions in nulliparous rats, and primiparity increased EVs whereby these subregions are the most, or nearly the most, important within the networks of primiparous rats, regardless of genotype (EVs ranging from 0.92 to 1 across the subregions; Figure 3G.

The **retrosplenial cortex** presented with heterogeneity of subregion importance across groups, with low EV standard deviations in nulliparous wildtype and primiparous hAPOEε4 rats, and higher EV standard deviations in primiparous wildtype and nulliparous hAPOEε4 rats (Figure 3A). hAPOEε4 reduced RSA EV, regardless of parity (Figure 3H). RSGc responded differently, with high EV in nulliparous wildtype rats and extremely low EV in nulliparous hAPOEε4 rats (Figure 3H). Compared to nulliparous wildtype rats, primiparous wildtype rats had reduced RSGc EV, and primiparous wildtype and primiparous hAPOEε4 rats had similar EVs (Figure 3H). RSGa also responded with high EV in nulliparous wildtype rats, which was greatly reduced in nulliparous hAPOEε4 rats (Figure 3H). Interestingly, primiparity in wildtype rats lowered RSGab EV, whereas primiparity in hAPOEε4 rats increased RSGab EV (Figure 3H).

### A2. Heterogeneity in the importance of subregions within neural networks varied across the brain.

Some regions showed high homogeneity in eigenvector centrality, which implicates subregions were similarly important within the neural network, whereas heterogeneity might indicate a shift towards specific subregions being more important within the neural network. For example, subregions within the ventral hippocampus had low eigenvector centrality in primiparous hAPOEe4 rats, and only new neurons in the ventral DG had high eigenvector centrality. This might be related to the reported inhibitory effects of new-born neurons on mature granule cells (Anacker et al., 2018; Ash et al., 2023). Alternatively, the disengagement of older neurons in the ventral hippocampus during memory retrieval could have led to reorganization of the ventral hippocampus network to prioritize engagement of new neurons. However, primiparous hAPOEe4 rats exhibited fewer new neurons and lower activation of these neurons compared to all other groups (Lee et al., 2024). Together with the findings of the present study, this might suggest that new neurons, though sparsely activated, contributed uniquely to the neural network of primiparous hAPOEe4 rats. New neurons might have been selectively recruited for specific roles during memory retrieval, potentially with increased efficiency or specialization, as compensation for their reduced overall activation in primiparous hAPOEe4 rats.

Another apparent example of homogeneity was seen within the retrosplenial cortex in nulliparous wildtype rats and primiparous hAPOEe4 rats – although the retrosplenial cortex subregions were homogenously important in the network of nulliparous wildtype rats and not important in primiparous hAPOEe4 rats. In contrast, primiparous wildtype rats presented with a gradient of eigenvector centrality across subregions, with eigenvector centrality being highest in the RSA. Although nulliparous hAPOEe4 rats presented with generally low eigenvector centrality across retrosplenial cortex subregions, it was especially low in both RSGc and RSGab. As previously mentioned, hypometabolism and dysregulated functional connectivity of the retrosplenial cortex predicted conversion from mild cognitive impairment to AD in humans (Terstege, Galea, et al., 2024). Moreover, studies in both humans and rodent models have shown dysregulated functional connectivity of the retrosplenial cortex correlates with impaired cognitive performance (Mayne et al., 2024; Terstege et al., 2022). In sum, these findings highlight the nuanced and region-specific effects of primiparity and hAPOEε4 genotype on neural network organization and underscore the relevance of examining patterns of region importance as well as its variability to better understand the functional implications.

**A3. Errors in the delayed win-shift radial arm maze**

As previously reported, hAPOEε4 rats made more across-phase and within-phase errors than wildtype rats (across-phase errors main effect of genotype: *F*(1,41)=4.886, *p*=0.033, partial η^2^=0.106; within-phase errors main effect of genotype: p=0.085; Figure A2) (Lee et al., 2024). Errors decreased across time (across-phase errors main effect of block*: F*(3,123)=3.923, *p*=0.010, partial η^2^=0.087; within-phase errors main effect of block: *F*(3,123)=2.525, *p*=0.061, partial η^2^=0.058; Figure A2).

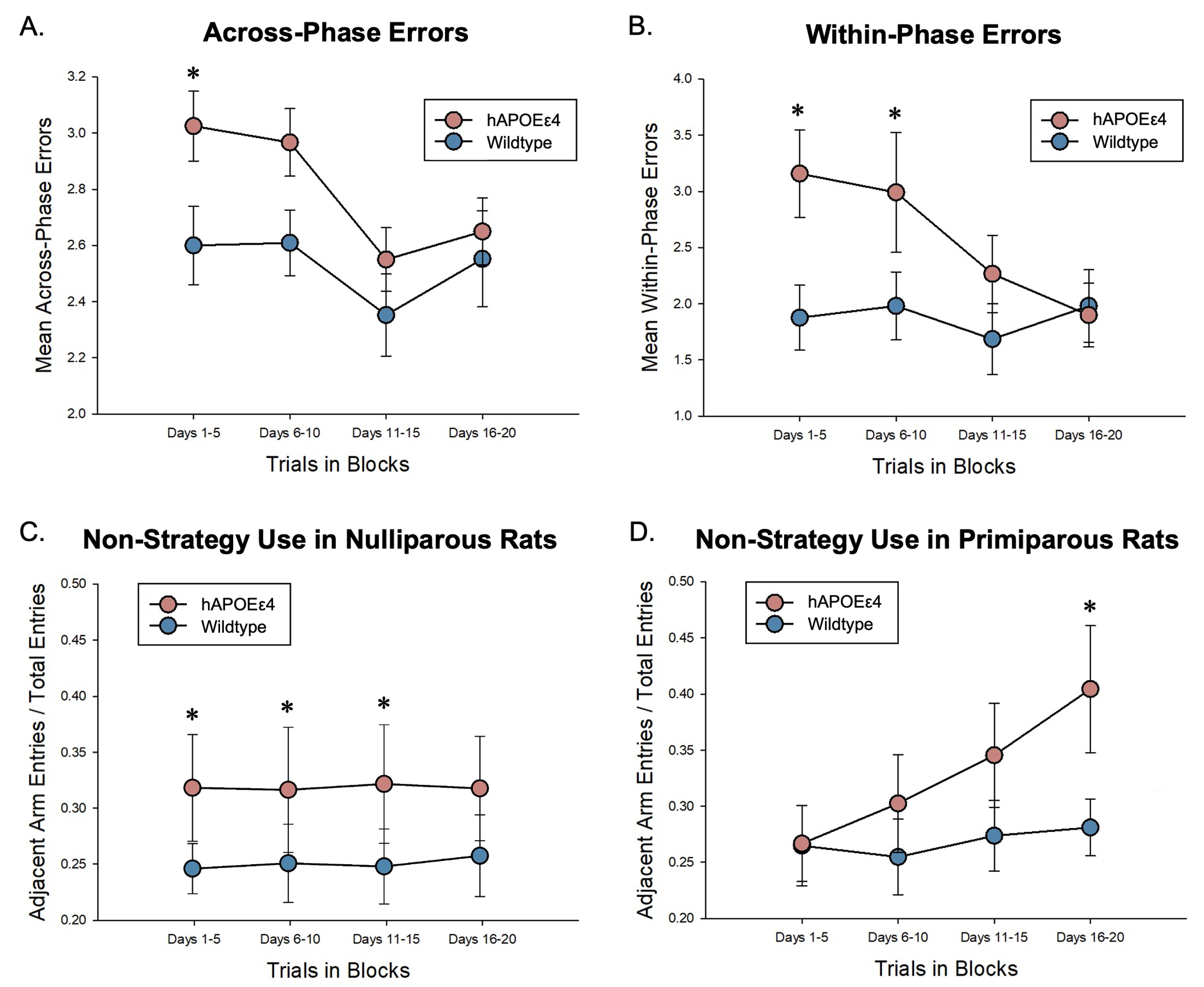

Figure A2. Mean number of across-phase errors (A) and within-phase errors (B) ± standard error of the mean in 5-day blocks in wildtype and hAPOEε4 rats. Reprinted with permission from (Lee et al., 2024).

### A4. Regions and combinations of regions for which neural activation was predictive of errors differed by parity and hAPOEε4 genotype.

We examined combinations of up to 4 regions to predict errors on the last day in the delayed win-shift radial arm maze task. For each group, the predictive power of each regional combination was assessed using adjusted R-squared values to account for the different numbers of predictors. Statistically significant region combination correlations with errors for each group were then ranked based on their adjusted R-squared values.

In nulliparous wildtype rats, subregions of the nucleus accumbens and dorsal striatum were involved in the strongest 2-region composites (NAc-NAs: adjusted R^2^=0.517, *F*=6.877, *p*=0.015; NAs-LA: adjusted r^2^=0.513, *F*=6.798, *p*=0.016; Figure 8A) and 3-region composites (vDG-mDS-RSGab: adjusted r^2^=0.707, *F*=9.846, *p*=0.005; vDG DCX-lDS-RSGab: adjusted r^2^=0.692, *F*=9.236, *p*=0.006; dDG DCX-NAc-RSGab: adjusted r^2^=0.640, *F*=7.520, *p*=0.010; Figure A3A) that were predictive of total errors. The strongest 4-region composite predictor of behavioural performance in nulliparous wildtype rats comprised of the mDS, hippocampal subregions, and hippocampal neurogenesis (dDG-vCA-vDG-mDS: adjusted r^2^=0.767, *F*=10.044, *p*=0.005; Figure A3A).

In primiparous wildtype rats, the strongest 2-region composite predictors of errors consistently included mDS, which was positively correlated with errors (mDS-lDS: adjusted r^2^=0.697, *F*=12.486, *p*=0.003; mDS-RSGab: adjusted r^2^=0.595, *F*=8.336, *p*=0.011; mDS-ACC: adjusted r^2^=0.534, *F*=6.734, *p*=0.019; Figure A3B). The strongest 3-region and 4-region composite predictors of behavioural performance involved subregions of the dorsal striatum and ACC (vCA3-NAc-ACC: adjusted r^2^=0.802, *F*=14.497, *p*=0.002; ACC-mDS-lDS: adjusted r^2^=0.798, *p*=0.002; ACC, mDS, lDS, and RSGC (adjusted r^2^=0.892; *F*=21.628, *p*=0.001; Figure A3B).

In nulliparous hAPOEε4 rats, the strongest region composite predictors of behavioural performance included more regions including subregions of the nucleus accumbens, hippocampus, and frontal cortex. The strongest 2-region composite predictors include: dDG DCX-ACC (adjusted r^2^=0.624, *F*=10.954, *p*=0.003), NAc-ACC (adjusted r^2^=0.559, *F*=8.600, *p*=0.007), and NAc-IL (adjusted r^2^=0.538, *F*=7.989, *p*=0.008; Figure A3C). The strongest 3-region composite predictors include: dDG DCX-ACC-NAc (adjusted r^2^=0.810, *F*=17.997, *p*<0.001), dDG DCX-ACC-RSA: adjusted r^2^=0.785, *F*=15.608, *p*=0.001; dDG DCX-ACC-NAs: adjusted r^2^=0.782, *F*=15.343, *p*=0.001; dDG DCX-ACC-IL: adjusted r^2^=0.734, *F*=12.051, *p*=0.002; Figure A3C). The strongest 4-region composite predictor of behavioural performance in nulliparous hAPOEε4 rats was made up of dDG DCX, ACC, vDG, and NAc (adjusted r^2^=0.882; *F*=23.509, *p*<0.001; Figure A3C).

In primiparous hAPOEε4 rats, the strongest 2-region composite predictors of errors included vCA3 and CeA, which were also the individual regions that significantly predicted errors (dCA3-CeA: adjusted r^2^=0.634, *F*=9.661, *p*=0.007; vCA3-vDG DCX: adjusted r^2^=0.521, *F*=6.432, *p*=0.022; IL-CeA: adjusted r^2^=0.521, *F*=6.429, *p*=0.022; vCA3-CeA: adjusted r^2^=0.503, *F*=6.068, *p*=0.025; Figure A3D). The strongest 3-region and 4-region composite predictors of errors included CeA and subregions of the hippocampus and retrosplenial cortex (vDG DCX-RSA-RSGab: adjusted r^2^=0.781, *F*=12.882, *p*=0.003; dCA1-vCA3-vDG DCX: adjusted r^2^=0.753, *F*=11.179, *p*=0.005; dCA1-vCA3-NAc: adjusted r^2^=0.745, *F*=10.721, *p*=0.005; dCA3-vDG DCX-CeA: adjusted r^2^=0.670, *F*=8.407, *p*=0.010; vCA3-vDG DCX-LA-RSA: adjusted r^2^=0.962; *F*=65.040, *p*<0.001 Figure A3D). The strongest 4-region composite predictor consisting of vCA3, vDG DCX, LA, and RSA predicted 96% of errors on the last day.

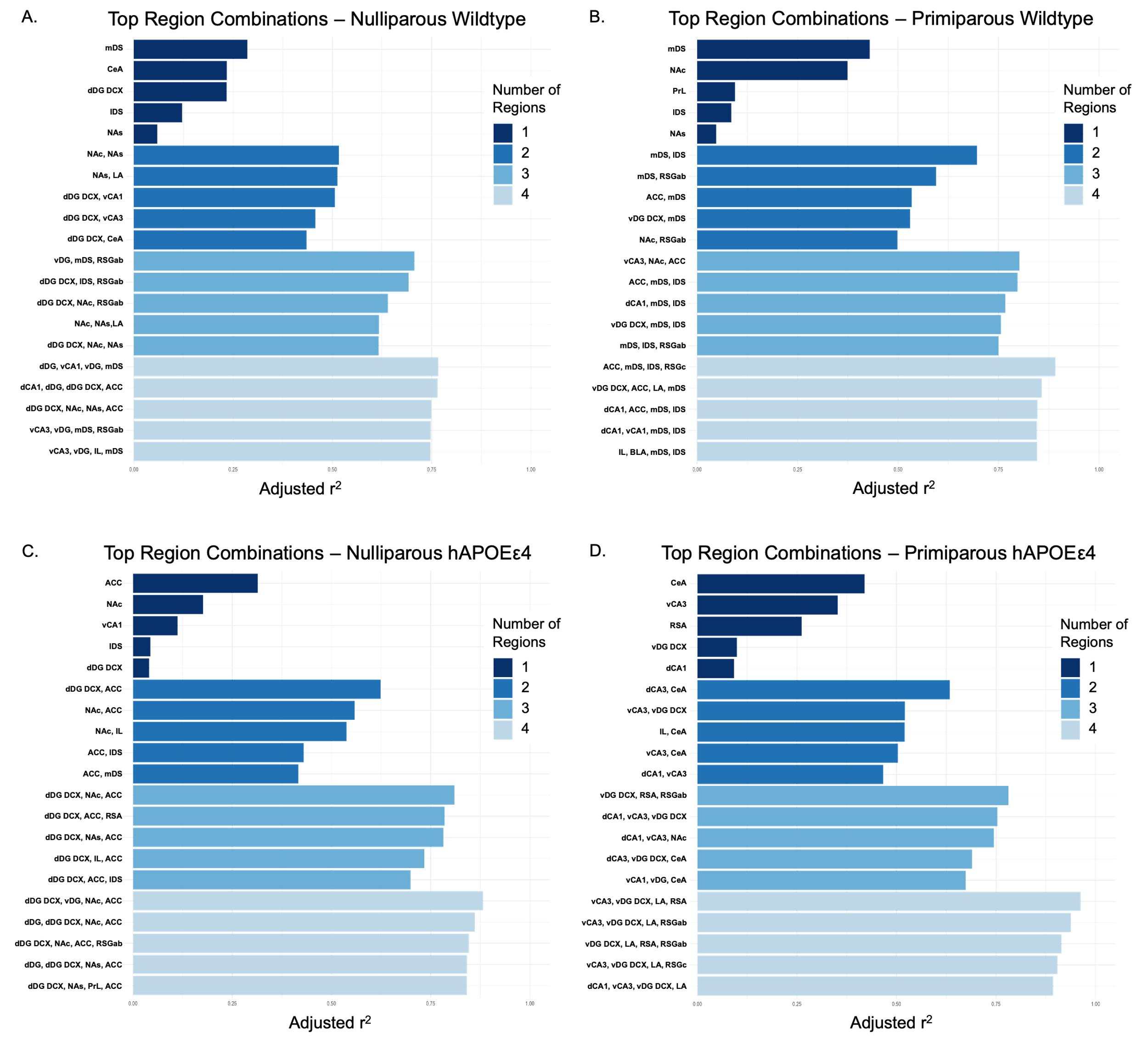

Figure A3. Multiple linear regression results for top regions significantly predictive of behaviour, which was operationalized as the total number of errors made on day 21 of the behavioural paradigm. CA – cornu ammonis, DG – dentate gyrus, DCX – doublecortin, IR – immunoreactive, NAc – nucleus accumbens core, NAs – nucleus accumbens shell, PrL – prelimbic cortex, IL – infralimbic cortex, ACC – anterior cingulate cortex, CeA – central amygdala, BLA – basolateral amygdala, LA – lateral amygdala, mDS – medial dorsal striatum, lDS – lateral dorsal striatum, RSA – agranular retrosplenial cortex, RSGc – granular retrosplenial cortex C (RSGc), RSGab – granular retrosplenial cortex AB, hAPOEε4 – humanized APOEε4
